# Measurement and comparison of acoustic space use in vocalizations of humans and close primate relatives

**DOI:** 10.64898/2026.06.14.732185

**Authors:** Hans T. Bilger, Michael J. Ryan, Julia A. Clarke

**Affiliations:** Department of Integrative Biology; University of Texas at Austin; Austin, TX, 78712, USA; Smithsonian Tropical Research Institute; Balboa, 0843-03092, Republic of Panama; Department of Geological Sciences; University of Texas at Austin; Austin, TX, 78712, USA

**Keywords:** Bioacoustics, evolution, language, signal evolution, speech, vocal production

## Abstract

The human larynx, compared to those of closely related primates, lies deeper in the throat and lacks vocal membranes and air sacs. These shifts are usually analyzed regarding their acoustic effects on vowel-like vocalizations, since the evolution of speech was long thought to require an expansion of vocal range driven by vocal tract modifications. However, vowels are just one type of phoneme, and speech is just one class of human utterance. To understand the evolutionary underpinnings of known shifts in human vocal morphology, a broader bioacoustic comparison is needed. Specifically, the range of sounds used in human speech must be compared to that employed in other human vocalizations and in the repertoires of extant close primate relatives. Here, we measure the acoustic-feature space occupied by human speech, non-linguistic, and musical vocalizations along with the calls of chimpanzees, bonobos, and chacma baboons. We use Mel-frequency cepstral coefficients to create an acoustic space depicting the spectro-temporal features of over 750,000 brief vocal segments sourced from published databases and other verified sources. Speech and song occupied significantly less volume in this acoustic space than human non-linguistic vocalizations. In addition, the acoustic-feature volumes of speech and song were not statistically distinct from those of non-human primates. These results suggest that speech was not enabled by an expansion of human vocal acoustic space. Anatomical shifts unique to humans may have led to an elaboration of non-linguistic utterances, but learned vocalizations use a surprisingly small fraction of this space. Our understanding of human vocal evolution will be further informed by additional systematic comparisons of the function and homology of non-speech vocalizations, along with the collection and incorporation of more complete non-human primate vocal datasets, especially from Gorilla and Orangutan.

## 1. Introduction

Complex vocalizations play an important role in human social structures, and can extend beyond the acoustic boundaries of speech and song to encompass a broader range of expressive sounds (Freeberg et al., 2012; McComb & Semple, 2005; Peckre et al., 2019; Sewall, 2015). Morphological changes in the vocal organ and tract that are unique to humans, relative to their nearest relatives, affect all vocal sounds produced (Fitch, 2000, 2018). However, complex human non-linguistic utterances produced with these modified structures have rarely been considered in comparative studies of human vocal evolution (Anikin et al., 2023).

The human vocal apparatus shows several derived morphological features compared to close primate relatives. First, the larynx lies lower in the throat and the tongue descends farther into the pharynx (D. Lieberman, 2011; P. H. Lieberman et al., 1969). Second, it lacks the supralaryngeal air sacs displayed by many monkeys and apes (De Boer, 2008; Hewitt et al., 2002). Finally, it has lost the thin, reed-like vocal membranes which lie above the vocal folds in all non-anthropoid primates (Nishimura et al., 2022). These shifts have often been linked with the evolutionary emergence of speech, which requires both the neuromotor ability to modify vocal output based on auditory experience—called vocal production learning—and a vocal apparatus mechanically capable of producing a sufficient number of perceptually contrasting sounds (Brenowitz & Beecher, 2023; Fitch, 2000; Janik & Knörnschild, 2021). Lowering of the larynx and lengthening of the tongue have been argued to increase the range of possible vowel-like articulations; air sac loss has been claimed to increase vowel discriminability; and the loss of vocal membranes (Fig. 1, A) has been proposed to stabilize the vocal source, making it simpler to produce a consistent bed of high-bandwidth tonal sound for subsequent vocal tract filtering (De Boer, 2012; Fitch, 2018; Fitch et al., 2016; Hewitt et al., 2002; P. H. Lieberman et al., 1969; Nishimura et al., 2022).

**Fig. 1.**
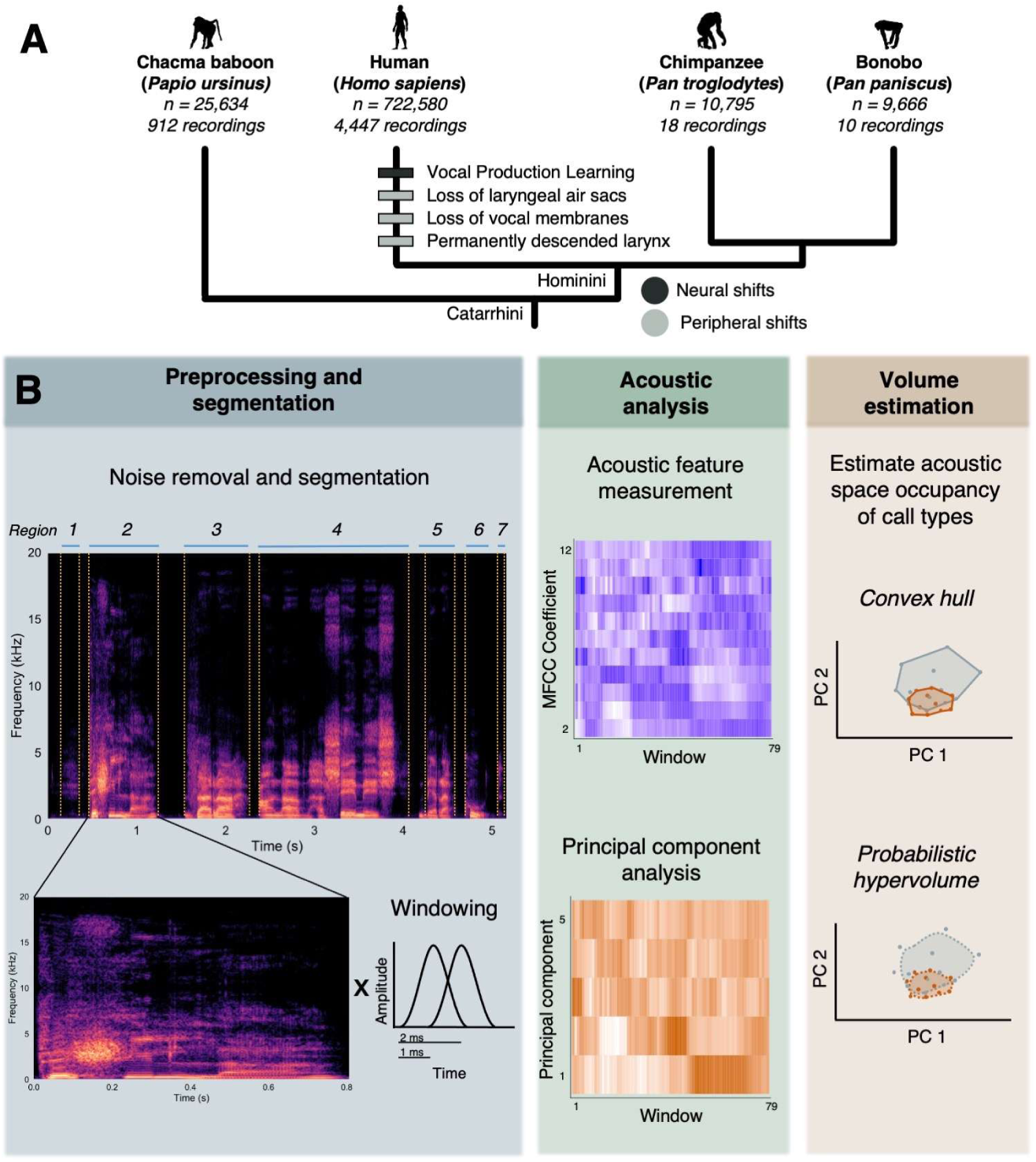
Evolutionary context for speech emergence and acoustic analysis pipeline. **(A)** Phylogenetic tree showing human autapomorphies relative to extant non-human primates. Shifts in the neural underpinnings of vocal control are marked in dark blue; peripheral shifts in vocal anatomy are marked in light gray. **(B)** Analysis pipeline for vocalization recordings. First (Preprocessing and segmentation panel), each recording was stripped of all possible extraneous noise and segmented into regions of quasi-continuous sound (top and bottom left). Then each region was split into a series of partially overlapping Hamming windows with a length of 0.02 s and an overlap of 50% (bottom right). Next (Acoustic analysis panel), each window was converted into 11 Mel-frequency cepstral coefficients. Principal component analysis (PCA) was then performed on this centered, scaled dataset of acoustic features. Finally (Volume estimation panel), the acoustic-feature space occupied by each vocalization type was estimated using convex hulls and probabilistic hypervolumes. PCs 1–5 were used to estimate these volumes. [Note to publisher: please print this figure in color.]

Speech-relevant shifts in the anatomy and neurology of the human vocal production system have led to distinct hypotheses of speech emergence. The “morphological” (or “peripheral”) hypothesis of speech emergence—that a morphology-driven expansion of human vocal output range enabled the evolution of speech—stands in contrast to the “neural” hypothesis of speech emergence, which ties vocal production learning (VPL) to speech origin (Darwin, 1871; Fitch, 2018). The morphological hypothesis was widely accepted for decades, but it has been challenged in recent years through studies demonstrating laryngeal descent in other taxa and the speech-capability of non-human primate vocal mechanisms (Boë et al., 2017; Fitch et al., 2016). It also remains untested in a crucial sense: Thus far, human vocal range shifts have mainly been measured using vowel-like articulations, which represent a small subset of our species’ vocal output.

Here, we assess evolutionary hypotheses of speech emergence by measuring the range of spectral envelopes, or “timbres”, present in a large portion of the human vocal output range and comparing it to those of key primate outgroups. To characterize these short-time envelopes, we used acoustic features known as mel-frequency cepstral coefficients (MFCCs). MFCCs provide a sparse, perceptually weighted numeric description of a vocalization’s short-time spectral envelope (Davis & Mermelstein, 1980). They were originally designed for human speech recognition tasks, and have since been used extensively in the study and classification of other primate vocalizations recorded in noisy and naturalistic environments (Cauzinille et al., 2024; Lakdari et al., 2024). While these features do not capture every aspect of vocal acoustic structure (such as fundamental frequency, syntactical arrangement, etc.), they have proven extremely useful for characterizing the various signatures of the upper vocal tract transfer function. The filter characteristic of the upper vocal tract is one of the chief perceptual contrast agents in human speech (it creates formant patterns and vowel identity), so the MFCC functions as a well-suited, perceptually-informed proxy of communication-relevant vocal range in humans and other primates.

Human non-linguistic utterances (cries, laughs, screams, etc.) are largely innate and produced through a broadly conserved neurological pathway older than and separate from the one used for generating speech and song (Jürgens, 2009). By comparing the volumes of the acoustic-feature spaces occupied by speech, song, non-linguistic vocalizations, and the vocal repertoires of extant primate relatives, we can empirically test for differences in the vocal output range of humans and non-human primates that lack human laryngeal novelties. We can also determine whether these differences are uniquely seen in learned vocalizations (speech, song). These analyses allow us to discriminate between alternative scenarios of speech emergence, since the morphological hypothesis predicts a speech-centered expansion of the range of human vocal sound qualities due to human-specific anatomical shifts, while the neural hypothesis does not. Specifically, if the acoustic-feature volume occupied by human speech sounds is larger than that of nonlinguistic vocalizations or the vocalizations of nonhuman primates, this would support the morphological hypothesis. If the acoustic-feature volume of speech is less than or equal to that of the other groups, it would falsify the morphological hypothesis and be consistent with the neural hypothesis, which makes no predictions regarding evolutionary shifts in acoustic-feature volume.

## 2. Materials and Methods

### 2.1 Participant and experimental model details

Our dataset of 11,691 short segments of recorded human and non-human primate (chacma baboon, *Papio ursinus*; chimpanzee, *Pan troglodytes*; and bonobo, *Pan paniscus*) vocalizations from 5,388 individual recordings (Fig. 1A; Table 1, Table 2) is a novel synthesis of human and non-human Hominidae data. All recordings were sourced from published databases and other verified sources; since the study did not recruit novel participants, an ethics committee review was unnecessary. Chimpanzees and bonobos were included in the dataset because they are the closest living relatives of humans. Chacma baboons were included because they possess the most complete acoustic dataset available for the vocal output of a non-human primate. All three non-human taxa lack the human novelties described above. We were unable to locate a sufficiently comprehensive set of Gorilla (*Gorilla gorilla*) or Orangutan (*Pongo pygmaeus*) vocal recordings in the published literature, so these taxa were not included in the analysis.

**Table 1.**
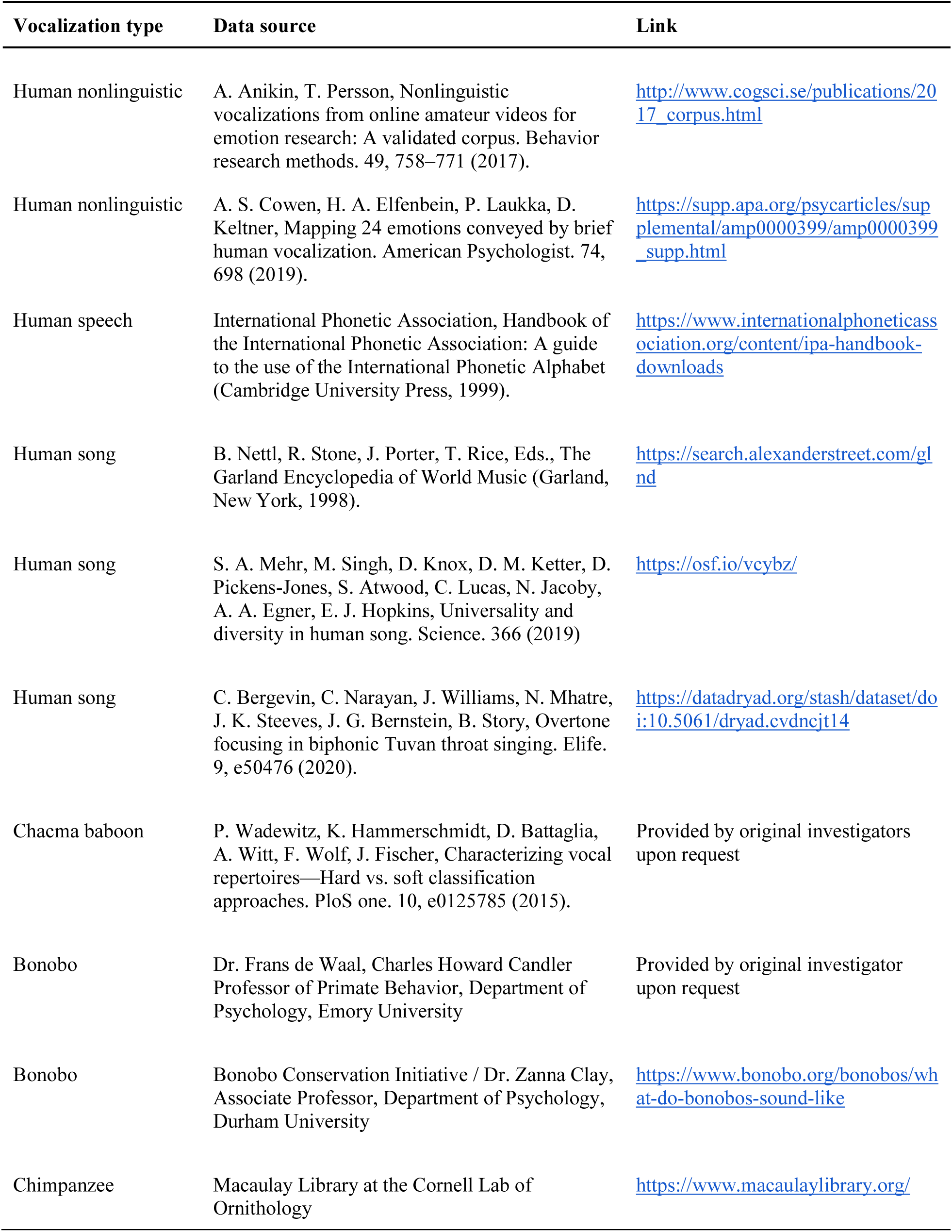
Recording sources.

**Table 2.** Acoustic data used in analyses of acoustic feature space. The table depicts the number of original recordings downloaded from each data source (“Recordings (*n*)”), the number of quasi-continuous vocal segments that each group of recordings was divided into following pre-processing (“Audio segments (*n*)”), and the number of partially overlapping Hamming windows (length = 0.02 s; overlap = 50%) that each group of audio segments was divided into (“Time windows (*n*)”).

| Vocalization type | Data source | Original sample rate | File type | Recordings (n) | Audio segments (n) | Time windows (n) | Learned vocalization? |
| --- | --- | --- | --- | --- | --- | --- | --- |
| Human nonlinguistic | Anikin and Persson 2017 | n/a | WAV | 563 | 1644 | 65691 | Some |
| Human nonlinguistic | Cowen et al. 2019 | n/a | mp3 | 2030 | 2030 | 199889 | Some |
| Human speech | International Phonetic Association 1999 | 20 kHz / 22.05 kHz / 44.1 kHz | WAV | 1660 | 3745 | 158953 | Yes |
| Human song | Nettl et al. 1998 | n/a | mp3 | 66 | 1387 | 221119 | Yes |
| Human song | Mehr et al. 2019 | n/a | mp3 | 118 | 354 | 44136 | Yes |
| Human song | Bergevin et al. 2020 | 96 kHz | WAV | 10 | 50 | 32792 | Yes |
| Chacma baboon | Wadewitz et al. 2015 | 44.1/48 kHz; later downsampled by original researcher to 20 kHz/5 kHz | WAV | 912 | 994 | 25634 | No |
| Bonobo | Dr. Frans de Waal | 44.1 kHz | WAV | 4 | 576 | 8203 | No |
| Bonobo | Bonobo Conservation Initiative | n/a | mp3 | 6 | 64 | 1463 | No |
| Chimpanzee | Macaulay Library at the Cornell Lab of Ornithology | 44.1 kHz | WAV | 19 | 847 | 10795 | No |

### 2.2 Recording sources

Human speech vocalizations were compiled by the International Phonetic Association and included a mixture of recordings of single words, phrases, and full sentences from adult male and/or female speakers. Recorded single words were chosen to represent the phonemic inventory of each language (International Phonetic Association, 1999). Twenty-seven languages from nine language families were represented in the corpus: American-English, Amharic, Arabic, Bulgarian, Cantonese, Catalan, Croatian, Czech, Dutch, French, Galician, German, Hausa, Hebrew, Hindi, Hungarian, Igbo, Irish, Japanese, Korean, Persian, Portuguese, Sindhi, Slovene, Swedish, Thai, and Turkish. No age or sex information was given for recordings except for those of Slovene, which contained vocalizations from male and female speakers.

Recordings of human singing were compiled from two broad cross-cultural samples of recorded human music mostly from small-scale societies (Mehr et al., 2019; Nettl et al., 1998) and a laboratory study of biphonic throat singing (Bergevin et al., 2020). In all musical corpora, only recordings of solo, unaccompanied singers were selected for further analysis. Human song can contain both linguistic and non-linguistic elements; all elements of song were included in the final dataset.

Human non-linguistic vocalizations were sourced from two databases. The first included 56 speakers (26 female, 30 male, ages 18–35, from USA, India, Kenya, and Singapore) who were asked to produce a variety of non-linguistic vocalizations relating to 30 emotional categories (Cowen et al., 2019). The second was an archive of 603 spontaneously produced human non-linguistic vocalizations, representing nine emotional categories, collected from YouTube (Anikin & Persson, 2017). In both cases, only vocal segments without obvious linguistic content were selected for further analysis.

Chacma baboon recordings were sourced from Wadewitz et al. (2015). This corpus comprised 912 recorded calls from wild chacma baboons in the Moremi Wildlife Reserve in Botswana. The dataset includes calls from 34 males, 35 females, 5 infant males, and 4 infant females. Recordings were selected to represent the species’ entire described vocal repertoire. Chimpanzee vocalizations were sourced from the Macaulay Library at the Cornell Lab of Ornithology. All recordings used in the final analysis were collected in the Mahale Mountains National Park and Gombe National Park in Tanzania. The recordings included a variety of call types from male and female adults and juveniles. Only recordings with a “five-star” rating that included mostly single callers were selected for further processing. The following recordings were used in the final analysis: ML 53994, ML 54074, ML 162001, ML 162110, ML 162158, ML 162306, ML 163515, ML 163593, ML 163596, ML 163604, ML 163634, ML 163639, ML 163641, ML 163650, ML 189988, ML 196196, ML 196248, ML 196251, and ML 196252.

Bonobo recordings were sourced from two locations. The first corpus of six recordings was provided by Dr. Zanna Clay and published by the Bonobo Conservation Initiative (Clay, 2022). This corpus included recorded calls of six types: “high hoots”, “food calls”, “travel calls”, “alarm and alert” calls, “threat barks and anger calls”, “male contest hoots,” and “laughter”. No information was provided on the age, sex, or location of callers. The second corpus was provided via personal communication by the late Dr. Frans de Waal. It included four digitized recordings of “hoots” and “screams” made by Bonobos in the San Diego Zoo. No information on caller age or sex was provided.

### 2.3 Bioacoustic analyses

#### 2.3.1 Recording pre-processing and segmentation

To measure the range of sound qualities contained in each vocalization category, we constructed a morphospace derived from acoustic-features known as Mel-frequency cepstral coefficients (MFCCs). MFCCs are human hearing-informed spectral summary features used for vocal signal analysis in a variety of taxa (Clink et al., 2019; Lee et al., 2006; Mielke & Zuberbühler, 2013). The MFCC was designed to characterize the short-term spectral envelope of human speech segments, so we assume it exaggerated variance in the human speech sounds in our dataset relative to other vocalizations. To prepare our audio data for MFCC extraction, each of the 5,388 recordings in the initial dataset was aurally and visually inspected. Recordings with high levels of environmental noise or obvious acoustic distortion were rejected. Pre-treatment noise attenuation has been shown to increase the performance of MFCC feature extraction (Soe Naing et al., 2020), so environmental noise and other acoustic artifacts were attenuated using a suite of audio processing tools in the program iZotope RX 9 Audio Editor (v9.2.0.1278; (iZotope, Inc., 2007)). Modules used within the program included “Voice De-noise”, “De-plosive”, and “De-clip”. Next, each recording was split into regions of quasi-continuous sound via visual assessment of spectro- and oscillograms (Fig. 1, B). For recordings with particularly high signal-to-noise ratios, the “Truncate Silence” feature in the audio recording and editing program Audacity (v2.4.2) was used to partially automate the segmentation process (Audacity Team, 2021). This process yielded 11,691 audio segments from <0.5 s to several seconds long. Next, each audio segment was converted to WAV format if necessary and resampled to 100 kHz using the *resamp* function in the R package *seewave* (Sueur et al., 2008).

#### 2.3.2 MFCC analysis

To calculate MFCCs, we first split each segment into a series of partially overlapping Hamming windows with a length of 0.02 s and an overlap of 50% (*n*=768,675). We then estimated the power spectrum of each window by conducting a short time Fourier transform and calculating its absolute value. This spectrum was then multiplied by a bank of frequency filters based on the Mel scale, a mathematical construct meant to approximate the response of the human auditory system. Finally, a discrete cosine transform was performed on the log of each filter bank’s total energy. This step has the effect of decorrelating and removing the vocal source signal (i.e. the pitch and harmonics of the voice) from the filter signal (i.e. broad spectrum shape and formant/vowel information), making it more robust at identifying speech-relevant acoustic features. Each of these transformed filter values constituted one Mel-frequency cepstral coefficient. We collected 12 MFCCs for each window within the range 0–20 kHz. No pre-emphasis filtering or liftering was used. MFCC features were collected using the function *melfcc* in the R package *tuner* (Ligges et al., 2018). Since the first MFCC for each time window chiefly describes the loudness of the segment rather than its spectral envelope, it was omitted it from subsequent analyses.

#### 2.3.3 Acoustic-feature volume analyses

Hypervolume analysis becomes extremely computationally expensive with increasing dimensionality, so principal component analysis (PCA) was performed on the centered, scaled MFCC dataset. Then, the first five PCs from each window (which accounted for ∼68% of cumulative variance) were used as input data to construct shapes for each vocalization type within acoustic feature space. The five-dimensional volumes of these shapes were then estimated using convex hulls and probabilistic hypervolumes (Blonder et al., 2018; Habel et al., 2019; Hutchinson, 1957).

Convex hulls were constructed using the “convhulln” function from the R package *geometry* (Habel et al., 2019; R Core Team, 2022). Hypervolumes were constructed using the “hypervolume_svm” function from the R package *hypervolume* (Blonder et al., 2022). The *hypervolume* package can use three different methods to construct hypervolumes: box kernel density estimation (“box”), Gaussian kernel density estimation (“gaussian”), and one-class support vector machines (“SVM”). We chose to use the “SVM” method because it responded most efficiently to the study’s large acoustic dataset. Default parameters were used for each application of the “hypervolume_svm” function.

Estimated acoustic-feature volumes were then compared across vocal types to test and generate hypotheses of acoustic space use in hominid vocal evolution. The patterning of points in acoustic-feature space offers information about the basic range of vocal sound qualities expressed by a given taxon or vocalization category, but says nothing about their higher-level temporal ordering (i.e., the rhythm or syntax of vocal phrases).

### 2.4 Quantification and statistical analysis

#### 2.4.1 Statistical validation—randomization tests

Results were statistically validated using randomization tests. For each comparison of interest (i.e. nonlinguistic vs. speech, human song vs. chacma baboon, etc.) group labels were shuffled and convex hulls/hypervolumes were recomputed. This process was repeated 10,000 times for each comparison (Figs. S1, S2). To determine whether the observed difference in hypervolume size was statistically significant, the following calculation was performed:

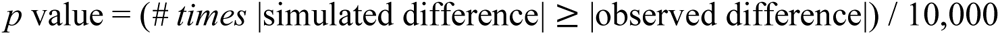

This yielded the *p*-values reported in Table S1. All statistical computations were conducted in R (R Core Team, 2022).

### 2.5 Data availability

#### 2.5.1 Materials availability

Information on the sources and accessibility of the original recordings analyzed in this manuscript is provided in Table 1. All recordings are available either in public repositories, for purchase from the original publisher, or through reasonable request to the respective original authors. Original recordings cannot be reproduced here because the authors of this manuscript do not have permission to mirror all original recording files used in the analysis.

#### 2.5.2 Data and code availability

Acoustic-feature data and example recordings have been deposited at Open Science Framework. A public DOI will be made available as of the date of publication. Data can currently be accessed at https://osf.io/e8vsr/?view_only=1badad869e2b439da63fc8e1139fdc68.

All original code has been deposited at Open Science Framework. A public DOI will be made available as of the date of publication. Code can currently be accessed at https://osf.io/e8vsr/?view_only=1badad869e2b439da63fc8e1139fdc68.

Any additional information required to reanalyze the data reported in this paper is available from the corresponding author upon request.

## 3. Results and Discussion

In both convex hull and hypervolume analyses, human speech was volumetrically smaller than non-linguistic vocalizations (randomization test, Fig. 2, A–C; Table 3). This result shows general agreement with the recent speech-nonverbal comparison carried out by Anikin et al. (2023), in which (1) nonverbal vocalizations showed significantly more fundamental frequency modulation than speech, and (2) speech sounds occupied a significantly larger amount of vowel space (as defined by the first two formant frequencies F1 and F2) than nonverbal sounds. The second result may initially appear contradictory with the present study, but here it is important to note that while vowel/formant space and MFCCs both capture aspects of the vocal filter function, they do not report exactly the same qualities. Formant frequencies are frequency ranges in the filter envelope that show local maxima, and they are determined solely by the configuration of the vocal tract (Fitch et al., 2025). MFCCs, on the other hand, report the relative acoustic energy present across all the frequency bands of the filter envelope—and the degree which those bands are excited depends partly on the behavior of the vocal source. For example, a low frequency scream-like vocalization and a spoken vowel could both use the same vocal tract configuration, resulting identical F1 and F2 frequencies. But in the case of the scream, high-frequency MFCC features would show greater values because of the added harmonics created by nonlinearities in the vocal source. Notably, Anikin et al. (2023) excluded two-thirds of screams and 10–25% of nonverbal vocalization types from their formant analysis since they were either “too high pitched for formant analysis or didn’t contain sufficiently long and stable vowel-like regions.” Taken together, the results of both studies suggest that while speech made richer use of the aspect of the vocal filter most important for lexical coding—vowel/formant space—it still reduced the number of general filter envelope shapes compared to nonlinguistic vocalizations.

**Fig. 2.**
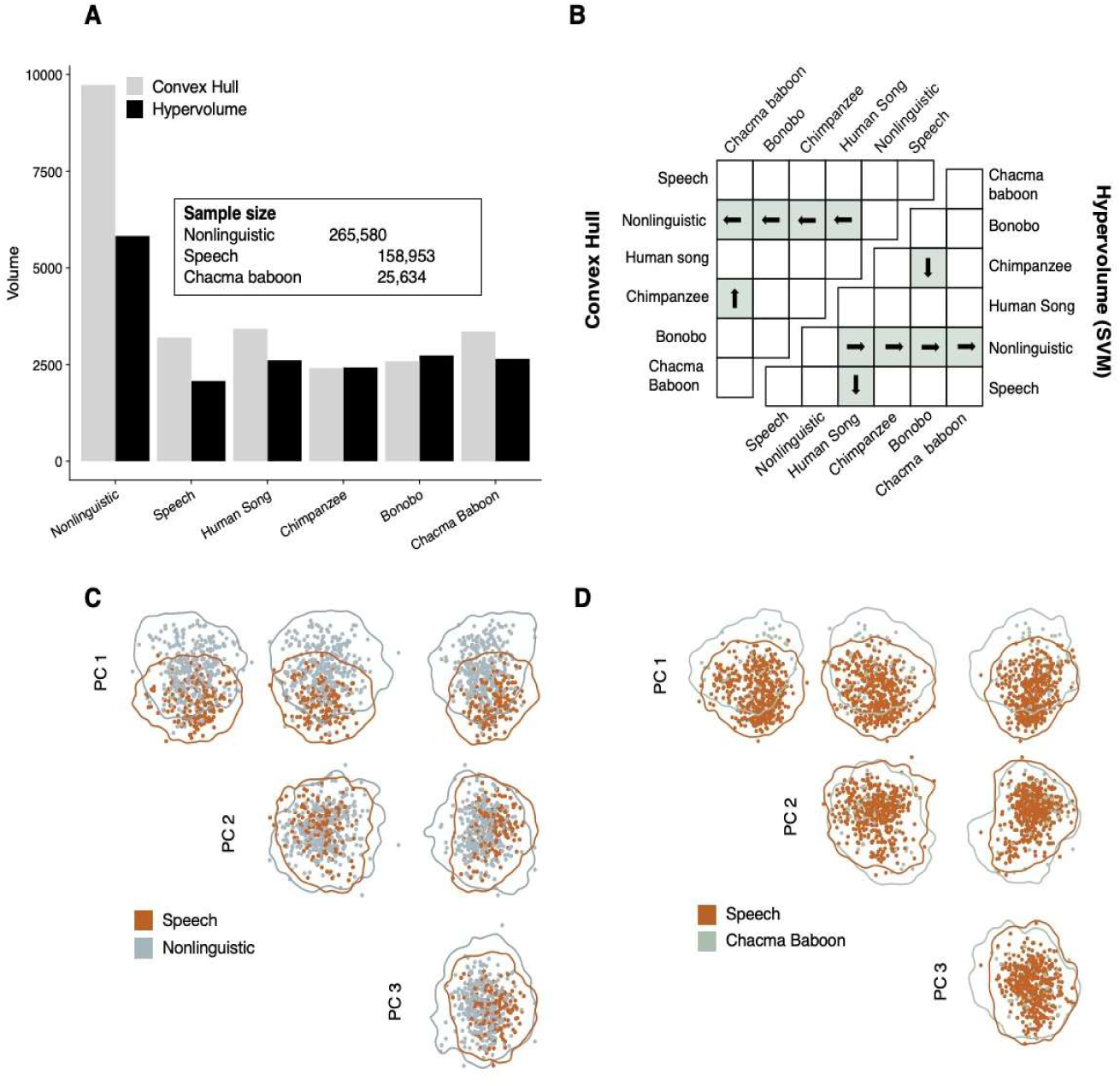
Non-linguistic vocalizations occupy more acoustic-feature space than speech, while speech and non-human primate outgroup repertoires occupy similar amounts. **(A)** Convex hull and probabilistic hypervolume volumes from a PCA including all data and constructed from MFCC 2–12. Non-linguistic sounds occupied more acoustic-feature space than speech, but the former group also contained more observations. Speech and chacma baboons occupied similar amounts of acoustic-feature space despite the former having more than 6x as many observations. **(B)** Differences in hypervolume size. Green color denotes significant differences; arrows point to the larger hypervolume in each dyad. Non-linguistic sounds occupied more acoustic-feature space than all other vocalization types in both analyses. No other significant group differences were recorded across both hypervolume and convex hull analyses. **(C)** Two-dimensional projections of hypervolumes for speech (orange) and non-linguistic vocalizations (blue). The larger size of the non-linguistic hypervolume is evident from the shape outlines. **(D)** Two-dimensional projections of hypervolumes for speech (orange) and chacma baboon vocalizations (green). The approximately equal size of the two hypervolumes is evident from the shape outlines. Chacma baboon was the non-human primate whose vocal repertoire was most fully captured in the dataset. [Note to publisher: please print this figure in color.]

**Table 3.** Observed acoustic-feature space volumes.

| Method | Group | Volume |
| --- | --- | --- |
| Hypervolume | Nonlinguistic | 5825.0 |
| Hypervolume | Speech | 2077.0 |
| Hypervolume | Human Song | 2611.5 |
| Hypervolume | Chimpanzee | 2429.0 |
| Hypervolume | Bonobo | 2734.7 |
| Hypervolume | Chacma Baboon | 2647.2 |
| Convex Hull | Nonlinguistic | 9727.7 |
| Convex Hull | Speech | 3200.9 |
| Convex Hull | Human Song | 3425.7 |
| Convex Hull | Chimpanzee | 2409.1 |
| Convex Hull | Bonobo | 2587.7 |
| Convex Hull | Chacma Baboon | 3359.8 |

The volumetric difference between speech and non-linguistic vocalizations can be partly explained through three of speech’s key features: It is quiet, informationally dense, and formant-centered. First, it is *quiet:* Unlike distress signals or advertisement calls, which are designed to propagate over large distances, speech is generally reserved for moderate-intensity, affiliative social interactions (Fitch, 2000). Some regions of acoustic space—roars, shrieks, hysterical laughter, etc.—can only be accessed at mechanical extremes of the vocal system. Such sounds could startle intended receivers or attract predators or eavesdroppers (Morton, 1977). Speech is also *informationally dense*. The limited temporal resolving power of the human auditory system along with the general tendency for biological systems to strive for maximally reliability at minimal metabolic cost (known as Zipf’s Principle of Least Effort) has pressured speech production to become more mechanically efficient over time (Coupé et al., 2019; Zipf, 1949). Many modern phonemes are “coarticulated”, meaning that their most extreme acoustic forms are traded for intermediate versions (Farnetani & Recasens, 2010). This simplifies the transitions between vocal tract configurations at the cost of reduced acoustic contrast between phonemic elements and increased load on the auditory brain.

The relatively low-contrast, highly informational quality of speech sounds partially explains why they are so *formant-centered.* Most vowel types are perceptually identified using the first two formant frequencies, and the spectral envelopes of these lower formants are easier to discern when they are excited by a low frequency source dense with harmonics (Anikin et al., 2023; Nishimura et al., 2022; Titze & Palaparthi, 2018). Furthermore, critical speech frequencies align with the region of the human auditory system with the best pitch discrimination. Average speech f0 is approximately 131 Hz for males and 220 Hz for females, with F1 frequencies of approximately 300–1,000 Hz (J. Hillenbrand et al., 1995; J. M. Hillenbrand & Clark, 2009). A strong sensation of “pitch” is only perceived for harmonic complex tones with a frequency range ∼30–4,000 Hz. Within that range, harmonic complex tones sound most “tonelike” (a feature associated with pitch and strongly associated with the formation of clear auditory images) between 82–784 Hz, with a maximum around 300 Hz (Huron, 2016; Oxenham, 2012). There is therefore a conspicuous alignment between frequencies used by the vocal production system to create speech-like sounds and the tuning of the auditory system that perceives them. Human screams can exceed 2 kHz in fundamental frequency (Pisanski et al., 2020), rendering that class of sounds less useful for the encoding of linguistic information.

The volume of acoustic space occupied by human speech was also and not significantly different from non-human primates in the present study (randomization tests, Fig. 2, A, B, D; Table 3). This result is—notwithstanding the caveats above regarding MFCCs and vowel space—consistent with recent work from Fitch et al. (2016) and Boë et al. (2017), who found non-significant differences in vowel space size between humans, macaques, and baboons. The lack of volumetric difference between speech and non-human primates was especially striking given disparities in sample sizes. Acoustic-feature volumes can only grow with increased sampling, and speech had ∼6x more observations than Chacma baboons, ∼16x more than chimpanzees, and ∼15x more than bonobos. These results are notable for the inclusiveness of the sampling. While prior comparative studies, including those of Fitch et al. (2016), have generally relied on single or more distantly related non-human primate species for their comparisons, the present study includes samples from the two closest human relatives (chimpanzee and bonobo) along with an example from a more distant outgroup (Chacma baboon).

Song and speech occupied similar volumes of acoustic-feature space (Fig. 2, A, B; Table 3). Song space was slightly larger in both convex hull and hypervolume analyses, but this difference was only significant in the latter. Since hypervolumes are less sensitive to outliers than convex hulls, our results indicate that the song data set contains a larger number of frequently used outliers than speech. Song is regularly transmitted over both long and short distances (consider opera singing vs. lullabies), so it might make more frequent use of acoustic extremes than speech. This quality would be in line with the finding that mammal and bird vocalizations with higher f_0_ radiate acoustic power more efficiently, making them more suitable for long distance communication (Titze & Palaparthi, 2018). This result is also roughly consistent with Anikin et al.’s (2023) analysis, which suggested that the range of fundamental frequencies produced in singing is greater than in speech and comparable with that of nonlinguistic vocalizations, with sex specific effects on the ranges of each vocal category.

Human song also figures in an influential hypothesis of language evolution known as the “musical protolanguage” (Darwin, 1871; Fitch, 2013). Since song requires vocal learning but not hierarchical syntax or complex semantics—two other key derived components of language—it has been proposed as an intermediate step between well-conserved innate calls and modern speech. This hypothesis suggests an evolutionary step where the sounds of the non-linguistic acoustic space were given a basic temporal ordering or coarse syntax. However, extant song often contains spoken elements, so it cannot be homologous with the “song” of the musical protolanguage—it is derived from speech. In our analysis, song occupied significantly less acoustic space than non-linguistic vocalizations and a similar amount to speech (randomization test, Fig. 2, A, B, Table 3). These results are consistent with extant song being speech-derived. It is notable that both vocally learned vocalizations in our dataset appear to have led to a contraction of the exploited acoustic space rather than the opposite. In each case, learning seems to have led to a refinement of timbral options rather than an expansion. A comparison with Anikin’s (2023) results is also informative here. Their analysis found that while both song and speech sustained particular fundamental frequencies for long, steady bouts, speech fundamental frequencies were lower in pitch and more restricted in range than singing fundamental frequencies.

Finally, the human non-linguistic space was significantly larger than that of all other vocalization types by at least a factor of two (randomization test, Fig. 2, A, B; Table 3). This could be partly due to innovation in the vocal periphery. The human loss of supralaryngeal air sacs and vocal membranes along with the acquisition of a permanently descended larynx and tongue have often been proposed to enable speech or make it more intelligible (De Boer, 2012; Nishimura et al., 2022). However, changes in the vocal organ and tract would affect all utterances. The first appearance of these shifts could equally have marked an increase in the diversity of human *non-linguistic* vocal sounds produced in new social contexts. An expanded non-linguistic repertoire, partially enabled by peripheral innovations and possibly concomitant with expansions in social-behavioral repertoire, may have preceded the evolutionary origin of spoken language. Social and communicative complexity have long been proposed to correlate, and vocal repertoire size appears to track with social bonding intensity within primates (Freeberg et al., 2012; McComb & Semple, 2005; Peckre et al., 2019; Sewall, 2015). This hypothesis requires further testing, however; the human non-linguistic space was better sampled than the non-human primate spaces in our study. This is especially true for bonobos and chimpanzees, which had a significantly smaller number of recordings in their respective samples.

Quantitative comparisons of vocal output have greatly informed the debate on speech origins, but prior work has focused on speech-like utterances. Exploration of the diversity of non-linguistic sounds in humans is relevant to the evolution of hominin vocal behavior.

Estimating vocal output range by collecting the broadest possible sample of vocal elements (Anikin et al., 2023; Odom et al., 2021) may lead to new areas of inquiry. We can better understand the evolution of vocal communication, including speech, by further embedding humans in comparative datasets that capture more inclusive samples of each taxon’s vocal output.

## Acknowledgments

The authors thank the late F. de Waal, The Macaulay Library at the Cornell Lab of Ornithology, The International Phonetic Association, the Bonobo Conservation Initiative, A. Anikin et al., A.S. Cohen et al., Bergevin et al., P. Savage, S. Mehr et al., and Z. Clay for supplying recordings; D. Spindler and T. Hess for statistical computing assistance; B. Blonder and D. Chen for guidance on probabilistic hypervolume analysis; A. Vickers for statistical guidance; and M. Knörnschild, B. Sealey, A.L. Baier, L. Legendre, and M. Hauser for support and comments. This work was completed in partial fulfillment of the requirements for a PhD at The University of Texas.

## Author contributions

Conceptualization: HTB

Methodology: HTB, JAC, MJR

Data curation: HTB

Formal analysis: HTB

Investigation: HTB, JAC, MJR

Visualization: HTB

Funding acquisition: HTB

Project administration: HTB

Writing – original draft: HTB

Writing – review & editing: JAC, MJR

Supervision: JAC, MJR

## Appendix A. Supplementary Material

Supplementary material to this article can be found online at [link].

